# A mechanism for attenuating responses to anticipated sounds in the dorsal cochlear nucleus

**DOI:** 10.64898/2026.08.22.746253

**Authors:** Qianyun Zhang, Salomon Z Muller, L.F. Abbott, Nathaniel B. Sawtell

## Abstract

The dorsal cochlear nucleus (DCN) is a cerebellum-like structure in the mammalian auditory brainstem that combines auditory nerve input with diverse auditory and non-auditory signals conveyed by granule cells. Granule cells form excitatory synapses onto inhibitory interneurons known as cartwheel cells, and *in vitro* studies have demonstrated an anti-Hebbian form of plasticity at these synapses. However, the function of cartwheel cells and their plastic granule cell input has remained unknown. Using *in vivo* electrophysiological recordings, optogenetics, and computational modeling, we provide evidence that intrinsic electrophysiological properties of cartwheel cells invert the expected effects of anti-Hebbian plasticity, generating a positive feedback loop that enhances cartwheel cell inhibition of DCN output neurons at the onset of anticipated sounds. This combined cellular and synaptic mechanism may implement a novel form of predictive processing that is robust to variability in the timing of anticipated sensory input.

## INTRODUCTION

The computations performed by the dorsal cochlear nucleus (DCN) have been enigmatic since the early descriptions of the histological structure of the mammalian cochlear nuclei by Lorente de Nó in the 1930s (De No 1933, Osen 1988). While both the ventral and dorsal divisions of the cochlear nuclei receive tonotopic input from the auditory nerve, the DCN resembles the overlying cerebellum in receiving an additional major source of excitation from granule cells (Mugnaini, Warr et al. 1980, Balakrishnan and Trussell 2008). Granule cells convey both auditory and non-auditory information, including somatosensory and vestibular signals (Oertel and Young 2004). The main targets of granule cell input are the cartwheel cells: a prominent class of interneurons that powerfully inhibit the output neurons of the DCN (**Figure 1A**) (Mancilla and Manis 2009, Roberts and Trussell 2010). Cartwheel cells share numerous similarities with cerebellar Purkinje cells, including firing two distinct types of action potentials: typical sodium spikes, called simple spikes and high-frequency calcium-dependent bursts, called complex spikes (Berrebi, Morgan et al. 1990, Zhang and Oertel 1993, Kim and Trussell 2007).

**Figure 1.**
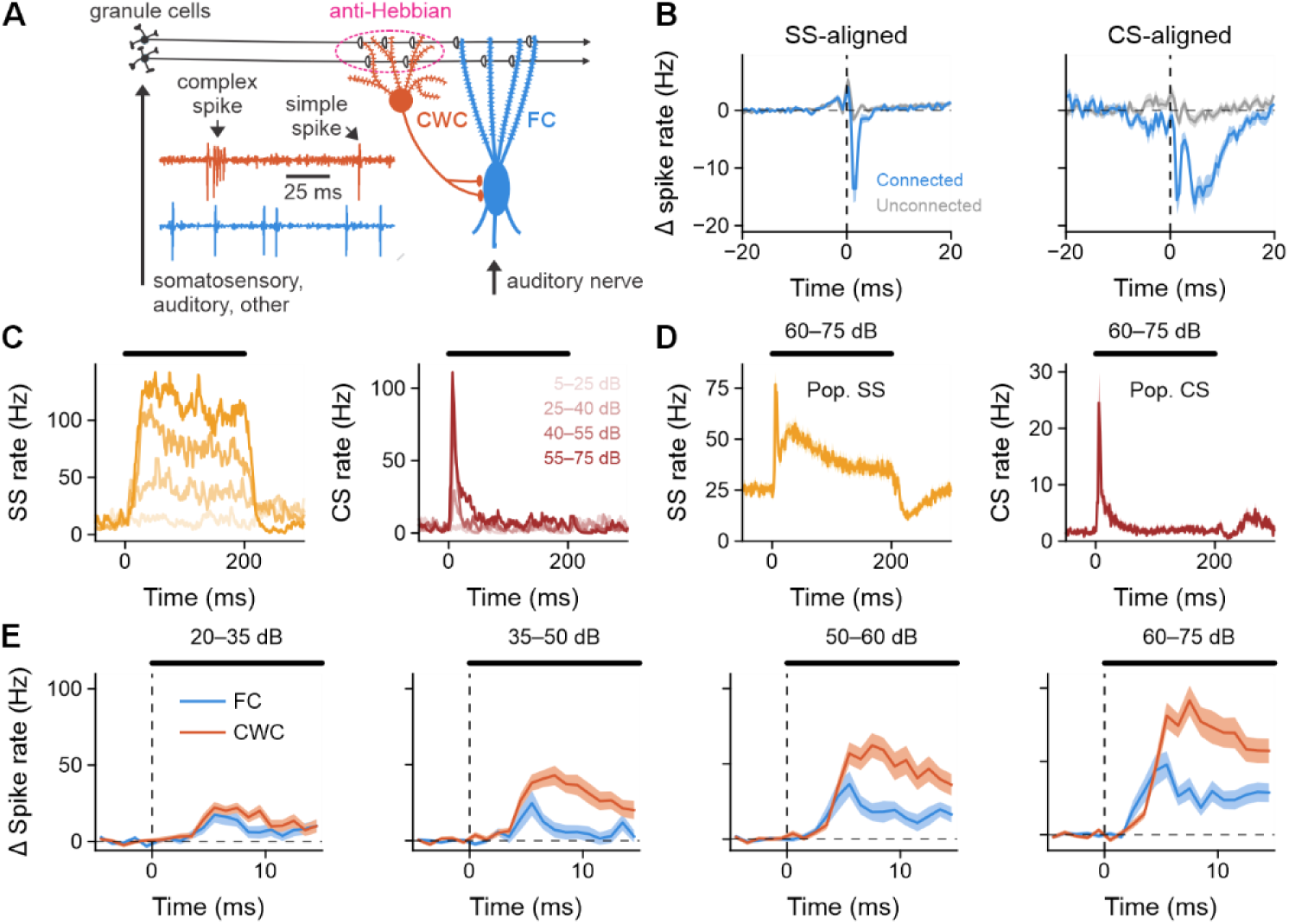
Temporal profiles of auditory responses in CWCs match FCs. **(A)** Schematic of DCN circuitry and example extracellular voltage traces from FC (blue) and CWC (orange) recordings. **(B)** Average cross-correlograms for putative monosynaptically connected (blue; n = 19) versus unconnected (gray; n = 50) pairs of simultaneously recorded CWCs and FCs. **(C)** Simple and complex spike responses to BBN across a range of sound levels for an example CWC. CS rate is calculated based on the first spike of each complex spike. **(D)** Population average simple and complex spike responses to BBN in CWC (n = 113). **(E)** Population average responses of FCs (n = 124) and CWCs (n =113) to the onset of BBN across sound levels. CWC spike rate includes simple spikes, complex spikes, and spikelets.

Complex spikes evoke large dendritic calcium influx and induce anti-Hebbian plasticity at granule cell synapses (Tzounopoulos, Kim et al. 2004, Roberts, Bender et al. 2008). In contrast to Purkinje cells, cartwheel cells do not receive input from the inferior olive and little is known about what drives complex spikes *in vivo* or their functional impact on DCN processing.

Among the functions proposed for the DCN is the attenuation of responses to self-generated, and hence partially predictable, sounds (Oertel and Young 2004, Bell, Han et al. 2008). Prior work showed that DCN output neuron responses to self-generated licking sounds are attenuated relative to responses to externally generated sounds and that silencing orofacial input to DCN granule cells reduces this attenuation (Singla, Dempsey et al. 2017). A well-studied form of sensory cancellation occurs in an evolutionarily related cerebellum-like structure in fish, the electrosensory lobe (ELL), where anti-Hebbian plasticity of granule cell input to Purkinje-like cells generates temporally-specific “negative images” that cancel responses to self-generated electrosensory input (Bell 1981, Bell, Han et al. 2008, Kennedy, Wayne et al. 2014, Muller, Zadina et al. 2019). This cancellation enhances the detection of behaviorally relevant external stimuli such as prey (Enikolopov, Abbott et al. 2018). Given the similarities in their circuitry (Bell, Han et al. 2008), could the DCN and ELL use similar cellular and synaptic mechanisms, including anti-Hebbian plasticity, to attenuate responses to predictable sensory input?

A problem with this hypothesis is that cancellation in the ELL relies on a precise temporal alignment between the negative image, which is computed based on motor corollary discharge signals conveyed by granule cells, and the sensory response it cancels. This alignment is possible due both to the physical features of electric fields and the extraordinary timing precision of the fish’s electrosensory and electromotor systems. In contrast, while self-generated sounds (e.g. those due to chewing) can be anticipated on the basis of orofacial sensory and motor signals, their exact timing (the moment a chip crunches, for example) can only be predicted approximately. In such a case, it is hard to imagine how a predicted “negative image” could be temporally-aligned with an actual chewing-related neural response with sufficient accuracy to cancel it. On the other hand, simply suppressing the system over an interval surrounding the time of an anticipated sound could significantly compromise its ongoing functions.

Here we provide evidence for a resolution to the problem of attenuating anticipated, but only partially predictable, signals. Our proposal, based on both experimental work and theoretical modeling, exploits parallels between DCN and ELL but relies on an essential difference: the presence of a T-type Ca^2+^ conductance in DCN cartwheel cells (Kim and Trussell 2007).

Remarkably, this conductance fundamentally changes the functional consequence of anti-Hebbian plasticity at granule-to-cartwheel cell synapses, despite the similarity of the underlying plasticity mechanisms to those in the ELL. In addition to providing a mechanism for attenuating anticipated sensory input, our analysis illustrates how the functional impact of a particular synaptic plasticity mechanism can depend critically on the biophysics of the connected neurons (Sjostrom, Rancz et al. 2008, Titley, Brunel et al. 2017, Magee and Grienberger 2020, Muller, Abbott et al. 2023).

## RESULTS

We characterized responses of cartwheel cells (CWCs) and DCN output neurons, known as fusiform cells (FCs), to external auditory stimulation (tones and broad band noise) and self-generated sounds during chewing behavior using acute multi-site extracellular recordings in awake head-fixed mice (Methods). Putative CWCs were identified based on the presence of distinctive high-frequency action potential bursts characteristic of complex spikes (**Figure 1A**) (Portfors and Roberts 2007, Ma and Brenowitz 2012, Singla, Dempsey et al. 2017). Putative FCs were identified based on their auditory response properties and high spontaneous firing rates (Young and Brownell 1976, Hancock and Voigt 2002, Ma and Brenowitz 2012). In a subset of simultaneously recorded units, cross-correlograms revealed putative monosynaptic inhibitory connections between CWCs and FCs (n = 19) and pairs of CWCs (n = 24) (**Figure 1B**). Complex spikes evoked a larger reduction in FC firing rate than simple spikes (**Figure 1B**). Furthermore, each spikelet in the complex spike response was associated with a similar-sized reduction in FC firing rate (**Figure S1**), consistent with prior *in vitro* evidence for faithful axonal propagation of high-frequency spike bursts in CWCs (Roberts, Bender et al. 2008).

To investigate how complex spikes may shape FC responses to sound, we examined the temporal profiles of responses to broad band noise (BBN) in CWCs (n = 108). Both simple and complex spike rates were strongly modulated by BBN in a majority of CWCs (∼92% and 83%, respectively). However, whereas simple spike firing typically remained elevated throughout the duration of the auditory stimulus, increases in complex spike firing were restricted to the onset, and in some cases, the offset of sound (**Figure 1C,D**). Unlike in Purkinje cells, where complex spike rates are negligible in comparison with those of simple spikes, complex spikes and their spikelets accounted for a substantial fraction of the peak firing rate response to BBN in CWCs (SS: 50.8 ± 35.4 Hz; CS: 16.5 ± 24.9 Hz; and CS + spikelets: 56.0 ± 85.6 Hz, mean ± sd). As expected from prior work, FCs recorded in the same sessions exhibited a variety of temporal firing patterns in response to BBN, including primary-like, buildup, and pauser (n = 116) (**Figure S2)** (Pfeiffer 1966, Godfrey, Kiang et al. 1975, Rhode, Smith et al. 1983, Parham, Bonaiuto et al. 2000).

To estimate the potential impact of CWCs on FC responses to sound we pooled responses across all recorded CWCs (including their simple spikes, complex spikes, and spikelets) and compared the temporal profile of this population response to that of FCs as a function of sound level (**Figure 1E**). Notably, the onset of CWC responses to BBN closely matches that of FCs, with CWC responses slightly lagging those of FCs but peaking at roughly the same time. Restricting analysis to simultaneously recorded CWCs and FCs with putative monosynaptic coupling yielded similar matching temporal profiles to the onset of BBN (**Figure S3)**.

### Complex spikes respond to transient features of sound

The surprising near simultaneity of CWC and FC responses raises the possibility that modifications in magnitude and/or timing of CWC onset responses could dramatically alter DCN output through CWC inhibition of FCs. To identify factors that could regulate the balance between auditory nerve excitation and CWC inhibition in FCs, we focused on complex spikes given their transient onset responses, their powerful inhibitory effects on FCs, and their role in inducing dendritic calcium influx and synaptic depression at granule cell synapses (Tzounopoulos, Kim et al. 2004, Roberts, Bender et al. 2008). *In vitro* studies have identified T-type calcium currents as key drivers of complex spikes in CWCs (Kim and Trussell 2007). Such currents require a period of membrane potential hyperpolarization for de-inactivation (Coulter, Huguenard et al. 1989). Consistent with this, we observed a clear decrease in simple spike firing preceding complex spike firing *in vivo* (**Figure 2A**). In the context of BBN stimulation, we found that the probability of evoking a complex spike (within 15 ms of sound onset) was higher when the rate of simple spike preceding sound onset (within a 60 ms window) was low (**Figure 2B**). Furthermore, plots of simple spike rate revealed a clear dip prior to trials on which a complex spike was evoked (**Figure 2C**). Similar dips in simple spike rate were observed prior to complex spikes evoked by the offset of BBN (**Figure 2D**) and the time of jaw opening during bouts of rhythmic chewing (**Figure 2E**). Together, these observations support a role for preceding hyperpolarization and T-type calcium currents in evoking complex spikes *in vivo*. Such a dependence on preceding hyperpolarization makes complex spikes well-suited to encode transient temporal features of sound and, as will be shown, has a major impact on the dynamics of synaptic plasticity in CWCs. Related roles for hyperpolarization-activated spike bursts in signaling input slope have been suggested for other sensory systems (Kepecs, Wang et al. 2002, Chacron, Longtin et al. 2004).

**Figure 2.**
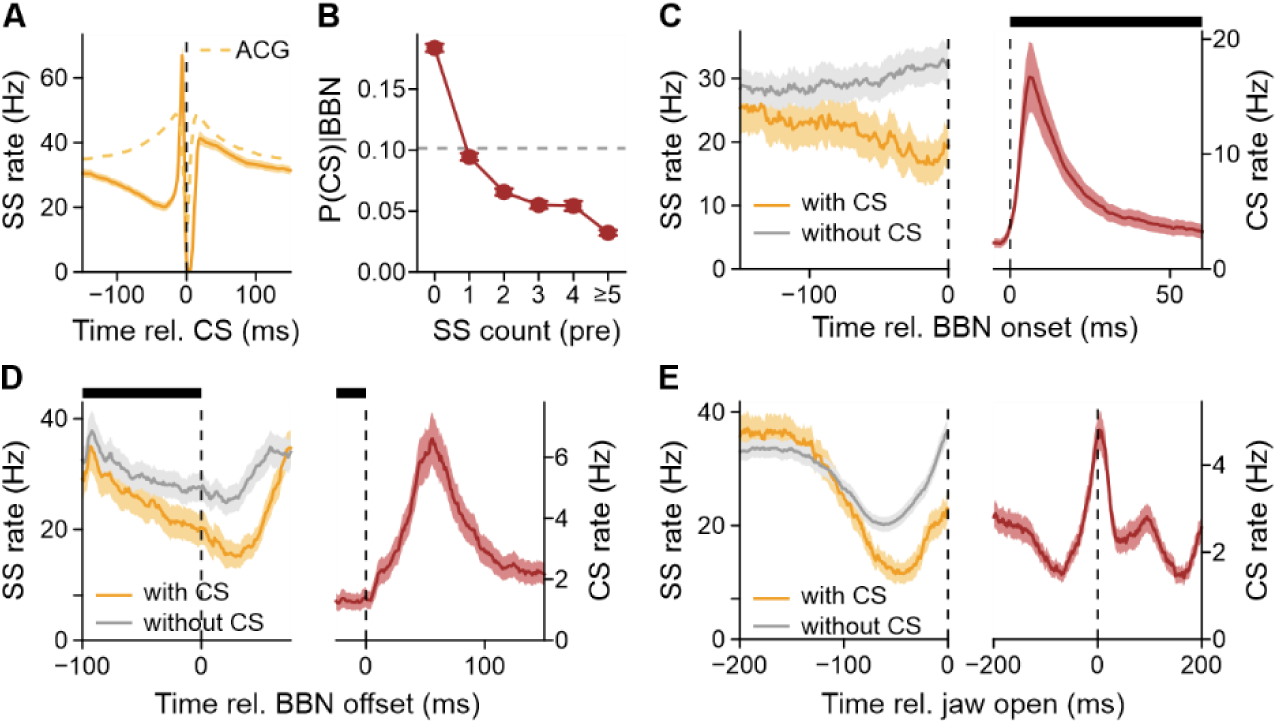
Factors promoting complex spike firing *in vivo*. **(A)** Average simple spike rate in CWCs aligned to the time of a complex spike (n = 114). Average simple spike auto-correlogram is shown for comparison (dashed). **(B)** Probability of BBN-evoked complex spiking depends on preceding simple spike rate (n = 109; >=40 dB SPL). **(C)** Simple spike firing rate aligned to BBN onset for trials with and without a complex spike (n = 67). **(D)** Simple spike firing rate aligned to BBN offset for trials with and without a complex spike (n = 65). **(E)** Simple spike firing rate aligned to jaw opening for trials with and without a complex spike (n = 106).

### *In vivo* plasticity of chewing-related input to CWCs

To probe plasticity, we optogenetically activated CWCs in closed-loop during rhythmic chewing behavior. Chewing is expected to engage somatosensory input to DCN granule cells from the spinal trigeminal nucleus (Zhou and Shore 2004, Singla, Dempsey et al. 2017, Balmer and Trussell 2021). Channelrhodopsin-2 expression in the DCN was restricted to CWCs using a Calb1-Cre driver line (**Figure 3A**; Methods). High-speed video was used to trigger optogenetic stimulation based on the tracked position of the jaw. Blue light pulses (10 ms) were delivered through an optical fiber integrated with the recording probe, allowing for simultaneous optogenetic activation and recording of CWCs as well as recording of nearby FCs (**Figure 3B,F**). To look for plastic changes, we computed the difference in CWC and FC responses during bouts of rhythmic chewing before and after a pairing period (∼15 minutes) in which light stimulation was delivered just following jaw closure (**Figure 3C**,**G**; dashed lines, after-before). We also computed the change in CWC and FC responses during the pairings (**Figure 3D,E,H**; late – early).

**Figure 3.**
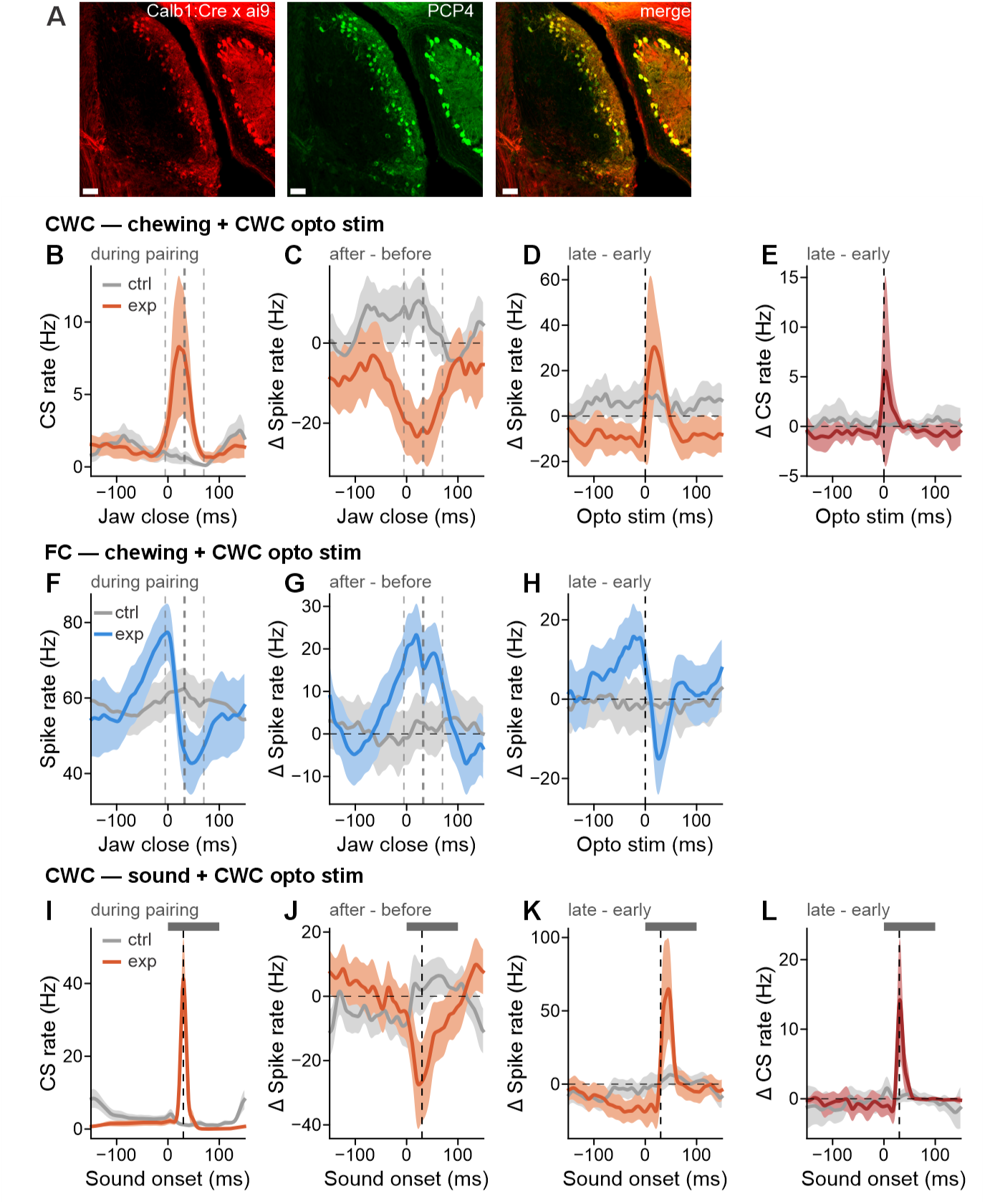
Dynamics of CWC plasticity *in vivo*. **(A)** Coronal section from a Calb1-Cre × Ai9 mouse immunostained for PCP4 (green). tdTomato (red) labels Calb1+ neurons; colocalization with PCP4 (merge, yellow) confirms selective targeting of CWCs within the DCN. Note, Purkinje cells are also labeled in adjacent cerebellar regions. Scale bars, 100 µm. **(B–E)** Effects of closed-loop optogenetic stimulation of CWCs during chewing on CWC firing (exp, orange: *n* = 9 cells, ctrl, gray: *n* = 4 cells, shading indicates SEM). Dashed vertical lines indicate mean ± SD of the timing of optogenetic stimulation pulses relative to jaw close (mean 32 ms, SD 38 ms). Complex spike (CS) rate aligned to jaw-close events during optogenetic stimulation **(B)**, pairing-induced change in total spike rate **(C)**, and change in total spike rate **(D)** and CS rate **(E)** during pairing aligned to the delivery of the optogenetic stimulation**. (F–H)** Effects of closed-loop optogenetic stimulation of CWCs during chewing on FC firing (exp, blue; n = 8 cells; ctrl, gray; n = 13 cells, shading indicates SEM). Firing rate aligned to jaw-close events during optogenetic stimulation **(F)**, pairing-induced change in spike rate **(G)**, and change in spike rate during pairing aligned to the delivery of the optogenetic stimulation **(H). (I–L)** Effects of optogenetic stimulation of CWCs paired with auditory stimulation on CWC firing (exp, orange; n = 5 cells; ctrl, gray; n = 10 cells, shading indicates SEM). Gray bar indicates sound duration. CS rate aligned to sound onset during optogenetic stimulation **(I)**, pairing-induced change in total spike rate **(J)**, and change in total spike rate **(K)** and CS rate **(L)** during pairing aligned to sound onset.

After pairing, we observed a reduction in CWC firing rate around the time of jaw closure (**Figure 3C**), consistent with anti-Hebbian synaptic depression of chewing-related granule cell input. During pairing, we observed a reduction in CWC firing rate preceding the optogenetic stimulation along with increases in total spiking and complex spike firing evoked by optogenetic stimulation (**Figure 3D,E**). In light of the results shown in **Figure 2**, we hypothesize that increased complex firing is the result of preceding membrane potential hyperpolarization due to anti-Hebbian synaptic depression of chewing-related granule cell input. The effects of pairing on simultaneously recorded FCs were opposite to those observed in CWCs, consistent with CWCs inhibiting nearby FCs. FCs exhibited a firing rate increase around the time of jaw closure after pairing (**Figure 3G**) as well as an increase in firing prior to jaw closure during pairing (**Figure 3H**). Minimal changes in either CWC or FC responses were observed in littermate control mice lacking opsin expression (**Figure 3**, gray lines).

Finally, we performed a separate series of experiments in which optogenetic stimulation of CWCs was paired with a preceding BBN (**Figure 3I**). The results were similar to those described above for chewing. After pairing, CWCs exhibited a reduction in noise-evoked firing (**Figure 3J**). During pairing, we observed a reduction in CWC firing rate preceding the optogenetic stimulation along with a prominent increase in total CWC spike rate (**Figure 3K**) and complex spike firing (**Figure 3L**) evoked by the optogenetic stimulation.

### Evidence for attenuation of responses to anticipated sound during chewing

Next, we analyzed responses of CWCs and FCs during bouts of rhythmic chewing. We used multi-unit activity (MUA) recorded in the deep layers of the DCN (the termination site of auditory nerve fibers) as a proxy for chewing-related auditory input. MUA sites were confirmed to be strongly auditory based on their short-latency responses to BBN. MUA peaked around the time of jaw closure, presumably related to grinding of the food pellets (**Figure 4A**). The rate of audible crunches detected by a microphone positioned near the face also peaked at jaw closure. Simple spike firing was positively correlated with MUA (**Figure 4A,B**), consistent with our findings that simple spikes strongly encode broad band acoustic stimuli (**Figure 1C,D**).

**Figure 4.**
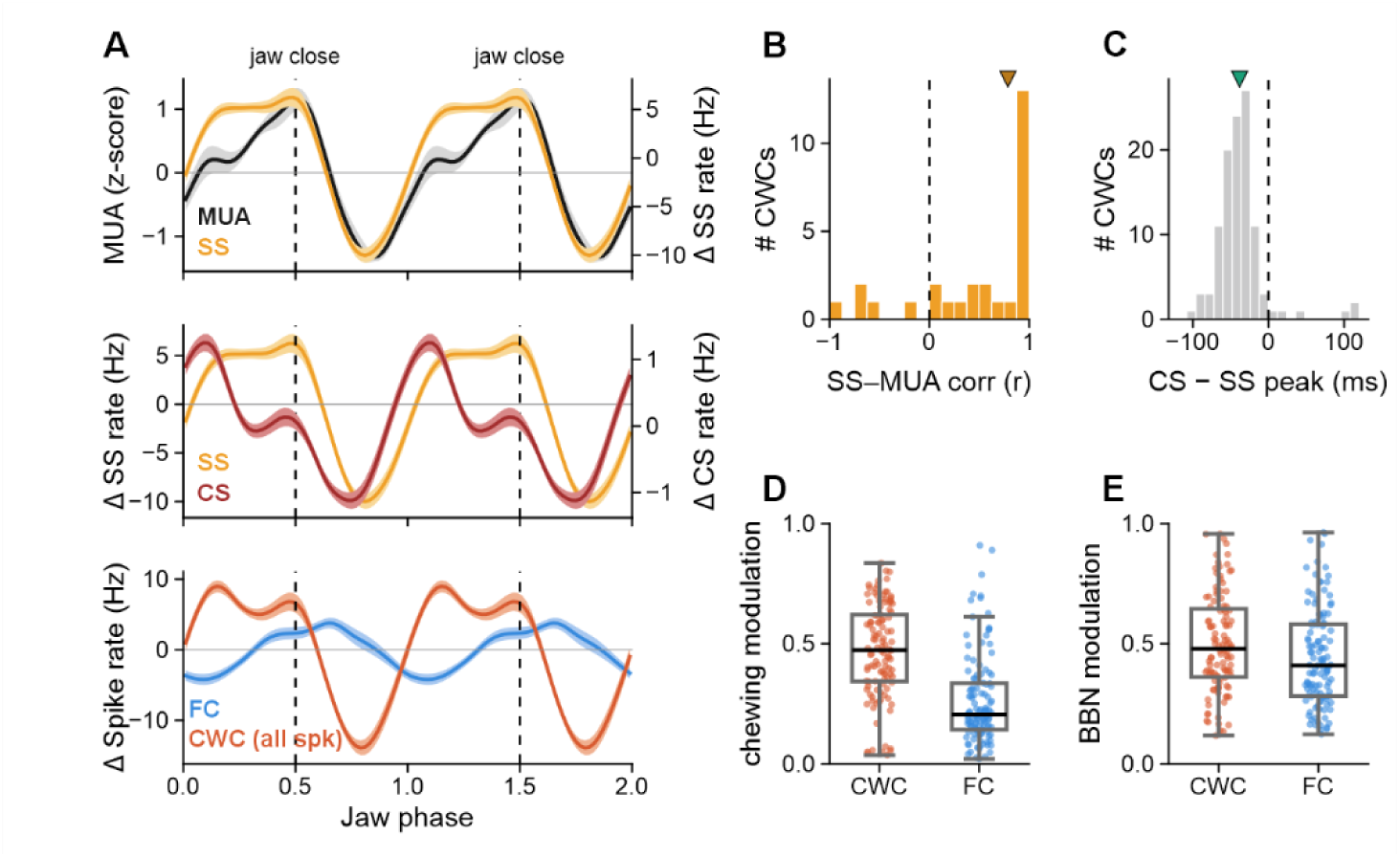
CWC and FC responses during chewing behavior. **(A)** Jaw-cycle phase averages (two cycles shown; dashed line = jaw closure at phase 0.5; mean ± SEM). Top: BBN-selected auditory multi-unit activity (MUA, z-scored; 14 sessions, black) with CWC simple-spike rate (SS, orange). Middle: CWC SS (orange) and complex-spike rate (CS, dark red). Bottom: fusiform-cell (FC, blue) and CWC all-spike (orange) rates (mean-subtracted). SS/all-spike, n = 114 CWCs; CS, n = 109; FC, n = 118. **(B)** Per-cell correlation between each CWC’s SS jaw-phase response and the simultaneously recorded auditory-MUA profile (n = 28 CWCs from the 14 MUA sessions; median r = +0.79), showing SS tracks the auditory input. **(C)** Per-cell CS − SS peak-time difference within the jaw cycle (n = 109; median = −38 ms, i.e., CS leads SS; Wilcoxon p = 3×10⁻¹⁵). **(D)** Chewing modulation index (MI = (max − min)/(max + min) of the jaw-phase rate responses), CWC total spike rate vs FC (n = 114, 118; Mann–Whitney p = 4×10⁻¹⁴). Boxplots: median and quartiles; points, individual cells. **(E)** BBN response modulation index (greatest peak or trough relative to pre-stimulus baseline; for the same cells (n = 112, 117; p = 0.03).

Complex spikes peaked earlier in the chewing phase than simple spikes (**Figure 4A,C**), consistent with evidence presented above that complex spikes encode transient features of sound (**Figure 1C,D**). Finally, we observed that CWCs were modulated much more strongly than FCs during chewing (**Figure 4A,D**). Responses to BBN were more similar across the two populations (**Figure 4E**), consistent with CWCs selectively attenuating FC responses to predictable sounds. Below we present a computational model of a cellular and synaptic mechanism that could mediate such an attenuation.

### A computational model of CWC function

To examine how previously described anti-Hebbian plasticity at granule cell synapses modifies CWC responses to anticipated sounds (Fujino and Oertel 2003, Tzounopoulos, Kim et al. 2004), we constructed a model CWC that receives input from a population of granule cells. The simple spikes rate in the model is given by a weighted sum of granule cell synaptic inputs, while the complex spike rate is a function of both the membrane potential (as reflected by the simple spike rate) and a T-type Ca^2+^ current (Methods). Auditory input to the model CWC was modeled as a total auditory granule cell input shaped to match CWC responses to BBN (**Figure 5A**, *red*). Signals predictive of this auditory input were modeled by a population of granule cell inputs with activity that tiles the time prior to and around the auditory input (**Figure 5A**, *gray*). In the case of chewing behavior, such signals could be provided by somatosensory mossy fiber inputs to granule cells from the spinal trigeminal nucleus (Zhou and Shore 2004, Singla, Dempsey et al. 2017, Balmer and Trussell 2021). As in our *in vivo* data, the increase in complex spike rate was limited to sound onset because the initial strong depolarization driven by the auditory granule cells inactivated the T-type channels thereby preventing sustained complex spike firing despite the sustained elevation in simple spike rate (**Figure 5B**) (Smith and Sherman 2002). Based on prior *in vitro* studies (Fujino and Oertel 2003, Tzounopoulos, Kim et al. 2004), plasticity was modeled with an anti-Hebbian plasticity rule in which granule cell inputs active prior to a complex spike are weakened, and those active without complex spike activity are modestly strengthened (**Figure 5C**, Methods). The precise shape of the anti-Hebbian learning rule was chosen to reproduce the temporal pattern of simple spike activity preceding complex spikes observed *in vivo* (**Figure 5D**).

**Figure 5.**
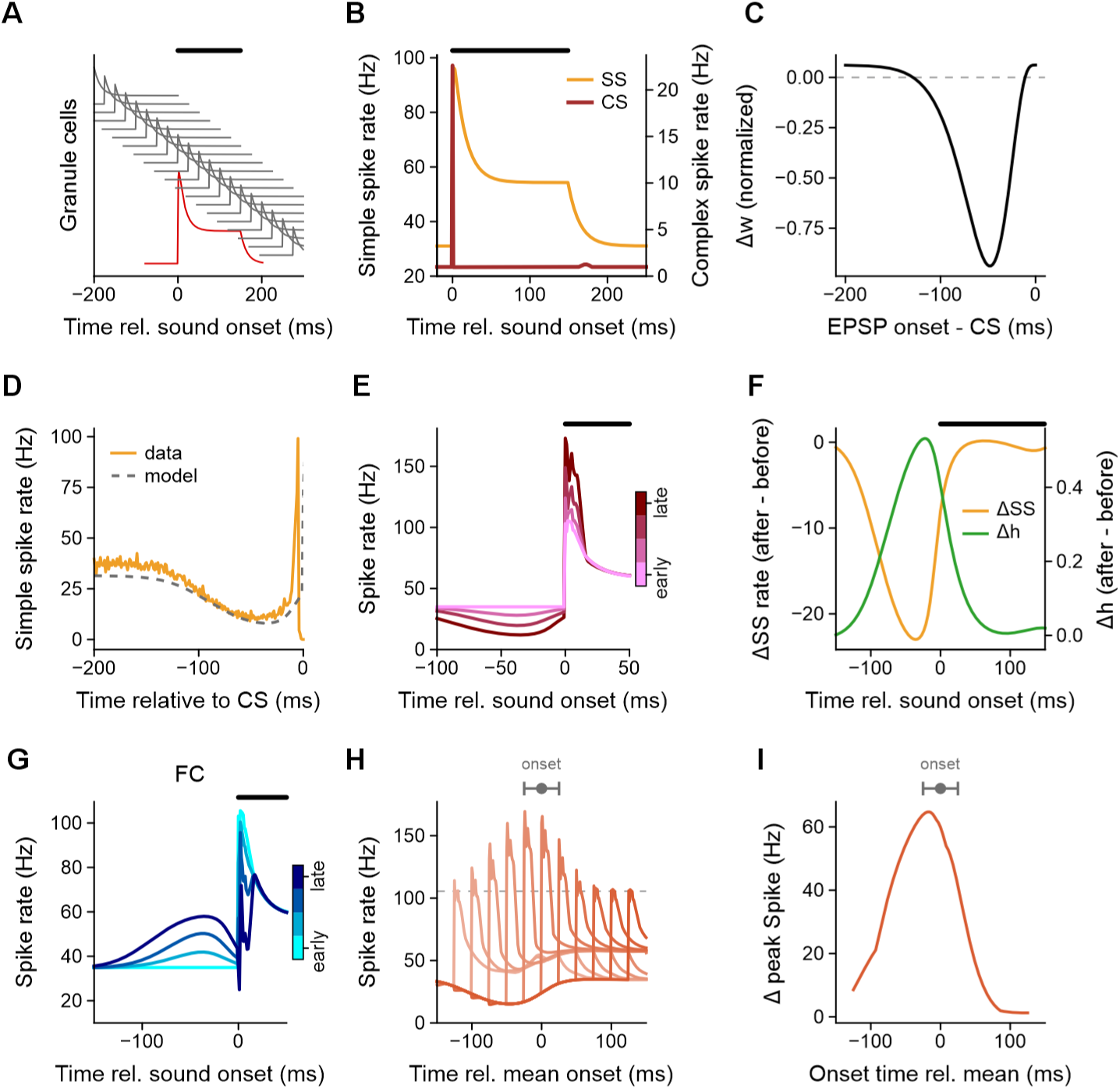
Simulations of CWC learning reveal a positive-feedback mechanism that enhances complex spike responses to predicted sound onset. **(A)** Schematic of model inputs to a CWC. Predictive granule cell inputs (grey) tile the time surrounding the sound, whereas auditory inputs (red) provide excitation beginning at sound onset. The model also includes inhibition simulating superficial stellate cells (not shown). **(B)** Model CWC response before learning. Auditory input drives a transient increase in simple spike (SS) and complex spike (CS) rates at sound onset, followed by a sustained increase in simple spike rate. **(C)** Anti-Hebbian plasticity rule used in this version of the model. The rule contains a broad depression window and a flat region with little weight change for granule cell inputs active immediately before a complex spike. **(D)** Simple spike activity aligned to complex spikes. The solid trace shows the average *in vivo* simple spike rate aligned to complex spikes occurring around jaw opening during chewing. The dashed trace shows the model simple spike rate aligned to sound-onset complex spikes after learning. The precise shape of the anti-Hebbian learning rule in **C** was chosen to reproduce the temporal pattern of simple spike activity preceding complex spikes observed *in vivo*. **(E)** Learning-induced changes in the model simple spike rate and availability of the modeled T-type current. Learning reduces simple spike firing before sound onset (orange), producing relative hyperpolarization that increases the T-type current, quantified by the change in the inactivation variable *h* (green). Traces show the difference between activity after and before learning. **(F)** Learning decreases CWC firing before the predicted sound and enhances the response at sound onset. Left, total CWC spike rate across learning, from early to late trials. **(G)** Effect of CWC learning on a model FC. FC firing is shown across learning, from early to late trials, illustrating the progressive reshaping of the response around predicted sound onset with a pre-sound ramp and attenuation of the onset response. **(H-I)** Learning-induced enhancement of CWC sound-onset responses is robust to variability in sound timing. **(H)** Total CWC spike rate for sounds occurring at different times relative to the mean predicted onset; the horizontal dashed line indicates the peak response to an unpredicted sound. **(I)** Change in peak CWC spike rate relative to the unpredicted-sound response as a function of the sound timing. Mean and ± std of the timing of the sound are shown above. Enhancement occurs across a broad temporal window corresponding to the period of increased T-type current availability.

A sound input to the model CWC triggers complex spikes at the time of the sound onset (0 in **Figure 5E**). When this sound is accompanied by predictive granule cell input, anti-Hebbian plasticity triggered by the complex spikes reduces the strength of the granule cell synapses carrying earlier predictive input. This causes a reduction in the rate of CWC simple spike firing prior to the time of the sound onset (**Figure 5E**, times < 0; the activity in **Figure 5E** shows the sum of simple spikes and spikelets due to complex spikes, as both of these contribute to inhibiting FCs). Normally, we would expect such a reduction in synaptic drive to reduce the probability of the sound onset triggering a complex spike. This, in turn, would reduce the plasticity until an equilibrium was reached, which is the process underlying negative image formation in the ELL. Here, however, the complex spike response is *enhanced* rather than reduced by the reduction of granule cell input. This is because the hyperpolarized membrane potential resulting from this plasticity (reflected in the pre-sound reduction of spiking in **Figure 5E**) de-inactivates T-type Ca^2+^ channels which enhances complex spike firing (**Figure 5F**). In other words, the hyperpolarization induced de-inactivation of the T-type Ca^2+^ conductance turns what would normally be a homeostatic process with negative feedback into a learning process with positive feedback. The result is to reduce overall CWC spiking prior to the sound onset and to enhance the transient CWC response to the sound.

In summary, the combination of anti-Hebbian plasticity and a hyperpolarization-activated inward current results in two modifications of the CWC response, as measured by the sum of simple spikes and complex-spike associated spikelets: 1) the transient response to the sound onset is enhanced and 2) the response decreases prior to the anticipated sound. Enhanced CWC inhibition at the time of the sound onset is expected to reduce and/or truncate the auditory response of the FC (**Figure 5G**). The functional impact of the reduction of inhibition prior to sound onset is less certain (see **Discussion**) but may serve to reduce the slope or “smooth” the FC response around the time of sound onset.

We have stressed from the outset that any attenuation mechanism must operate even when the times of sound onset cannot be predicted precisely. To test our model in this regard, we simulated a scenario in which the timing of the sound relative to the predictive input was randomly drawn from a normal distribution with a 25 ms standard deviation (**Figure 5H**). In this scenario, plasticity drove an increase in CWC responses to sound across a range of temporal offsets relative to the mean delay (**Figure 5H,I**). As long as sound-onset occurred within the temporal window in which the T-type current is elevated, the complex spike response increased.

For the model we have described to work properly, granule cell inputs that predict an upcoming sound should be weakened by synaptic depression but, because the mechanism relies on a strong and rapid CWC sound response, synapses carrying the sound input should not be depressed. We identified two ways to assure that this does not happen. One is to utilize a temporal window for anti-Hebbian plasticity that reproduces features of our *in vivo* data but which is slightly wider and delayed relative to that measured *in vitro* (**Figure 5D**). With this rule, inputs immediately preceding the sound response (with a latency less than this delay) are not depressed. Another option is to use the temporal window measured *in vitro* but to assume that the subset of granule cell conveying auditory input are non-plastic (**Figure S4**). Anatomical evidence for distinct sets of granule cell input to CWCs is discussed below. Finally, our model also assumes that granule cells conveying predictive inputs are not sufficient to evoke complex spikes on their own. This requirement could be met if granule cell excitation was approximately balanced by inhibition. Consistent with this, the DCN molecular layer contains interneurons (known as superficial stellate cells) that receive granule cell input and inhibit CWCs and FCs (Apostolides and Trussell 2014).

## DISCUSSION

This study synthesizes new findings with a large body of prior work on the DCN to identify a combined cellular/synaptic mechanism for attenuating responses to the onset of anticipated sounds. Below we discuss important remaining questions regarding the neural implementation of the proposed mechanism, the possible significance of our findings for hearing, and how variations across convergently evolved cerebellum-like structures may support predictive coding strategies adapted to different sensory modalities.

### Origins of auditory responses in CWCs

While it has often been assumed that CWCs primarily process non-auditory signals conveyed by granule cells (Young, Nelken et al. 1995, Kanold and Young 2001, Oertel and Young 2004), our results demonstrate prominent CWC responses to broad band acoustic stimuli with temporal profiles similar to FCs (**Figure 1E**). The only prior study of CWCs in awake mice reported strong responses of CWCs to tones that matched nearby FCs in terms of their preferred frequency (Portfors and Roberts 2007). Prior studies in cats also reported strong auditory responses in a subset of CWCs (Parham and Kim 1995). Given that CWCs lack direct auditory nerve fiber input, the origins of the prominent, short-latency auditory responses observed here, and in prior studies, are a key question. Granule cells provide the main source of excitatory glutamatergic input to CWCs and are located within several separate domains within and surrounding the cochlear nucleus complex (Mugnaini, Warr et al. 1980). Granule cells located in an external shell surrounding the cochlear nucleus receive diverse non-auditory input, including somatosensory and vestibular input (Weinberg and Rustioni 1987, Burian and Gstoettner 1988, Ohlrogge, Doucet et al. 2001, Balmer and Trussell 2021), as well as nonprimary auditory inputs from the inferior colliculus, ventral cochlear nucleus, superior olivary complex, and type II auditory nerve fibers (Brown, Berglund et al. 1988, Caicedo and Herbert 1993, Oertel and Young 2004, Balmer and Trussell 2022). Notably, granule cells are also numerous in the deep layers of the DCN where primary auditory nerve fibers terminate (Mugnaini, Warr et al. 1980) and electron microscopy, Golgi, and axon degeneration studies in cat suggest that granule cells are among the direct targets of primary auditory nerve fiber terminals (Cohen, Brawer et al. 1972, Kane 1974).

Such a connection, if it exists in mice, could explain key features of our results and also suggests an intriguing parallel with the cerebellum where the ascending branch of the granule cell axon synapses locally onto Purkinje cells before bifurcating in the molecular layer to form PFs (Bower and Woolston 1983, Gundappa-Sulur, De Schutter et al. 1999). An analogous organization in the DCN would provide the basis for a functionally important division of labor in our model between auditory and predictive granule cell inputs (**Figure 5A**). Auditory granule cell inputs that trigger complex spikes at sound onset must be partially protected from depression: if these inputs were to undergo anti-Hebbian LTD, the positive feedback loop would be self-extinguishing rather than self-amplifying. Ascending segment contacts, if subject to different plasticity rules than parallel fiber synapses — as has been suggested in the cerebellum (Sims and Hartell 2006, Conti and Auger 2024) — could provide the fast, strong, stable auditory drive needed to sustain the loop, while weaker and slower (but more numerous) parallel fiber inputs carry rich predictive information that undergoes depression. Aspects of this hypothesis can be directly tested with connectomics.

### Impact of plasticity at granule cell to fusiform cell synapses

*In vitro* studies have shown that plasticity at granule-FC synapses follows a Hebbian spike-timing-dependent plasticity rule (STDP), in which elevated FC spiking drives potentiation of granule cell input that precede the increase in firing and depression of the granule cell inputs that follow it (Tzounopoulos, Kim et al. 2004). This rule is opposite in sign to the learning rule at granule cell-CWC synapses and, at first glance, would seem to oppose the learning that occurs through CWCs. Our results indicate that the effects of these two forms of plasticity are not, in fact, in opposition because the T-type Ca^2+^ conductance makes the CWC’s anti-Hebbian plasticity act in a pseudo-Hebbian manner. Thus, the CWC and FC plasticity may act in concert despite their apparent differences. Understanding the function of STDP in the DCN will also require taking the intrinsic biophysical properties of FCs into account, including transient potassium currents that regulate the temporal profile of FC auditory responses (Kanold and Manis 1999).

### Implications for auditory processing and behavior

Both the magnitude and temporal dynamics of responses to the onset of acoustic stimuli strongly influence auditory perception and behavior (Blumenthal and Keith Berg 1986, Heil 2003, Lorenzi 2007). By increasing the magnitude and decreasing the latency of CWC responses, anti-Hebbian plasticity could reduce behavioral responses to sounds that are partially predictable based on granule cell input. Anti-Hebbian plasticity also reduces CWC inhibition prior to the onset of an anticipated sound. A corresponding increase in the pre-sound response of FCs could effectively “smooth” their response to an anticipated sound (**Figure 5G**). DCN lesions in cat selectively disrupt reflexive orienting to acoustic stimuli (Sutherland, Glendenning et al. 1998, May 2000). Hence a prediction of the present work is that optogenetic activation of CWCs should reduce acoustic orienting. Such experiments are complicated at present, however, by off-target effects on Purkinje cells in the nearby cerebellar flocculus, which is labeled by the Calb1-Cre driver line used here and is itself implicated in head orienting behavior. Additional electrophysiological experiments in behaving mice are also needed to identify natural conditions under which the CWC plasticity mechanism described here can be studied.

### Comparative insights into circuit function

Similarities between vertebrate cerebellum-like structures, including the presence of anti-Hebbian plasticity at granule cell synapses onto Purkinje or Purkinje-like cells that fire two distinct types of action potentials, suggest that these structures perform related functions (Oertel and Young 2004, Bell, Han et al. 2008). However, these structures also exhibit striking differences in their connectivity, cellular properties, and plasticity rules. For example, complex spikes are generated by powerful climbing fiber input to Purkinje cells, backpropagating axonal spikes regulated by inhibitory and dis-inhibitory electrosensory pathways in the ELL of mormyrid fish (Muller, Abbott et al. 2023, Perks, Petkova et al. 2025), and hyperpolarization preceding strong auditory input in the DCN (Kim and Trussell 2007). The present study suggests that these differences may reflect adaptations supporting distinct forms of predictive sensory processing. In the ELL, anti-Hebbian plasticity provides negative feedback that cancels neuronal responses to sensory inputs by shaping granule cell input (e.g. motor corollary discharge) into “negative images” of predictable components of the sensory input. Remarkably, intrinsic biophysical properties of CWCs effectively flip the sign of this plasticity, allowing the predictable auditory input itself to elicit faster and stronger CWC-mediated inhibition of output neuron responses to sound. While the latter mechanism cannot achieve a precise cancellation of sensory input, as has been demonstrated for the ELL, it is nevertheless well-suited to provide selective attenuation of anticipated auditory inputs even when their timing is only partially predictable by granule cell inputs (**Figure 5H,I**). This may be a decisive advantage in the auditory system where numerous factors, including the complex, environment-dependent acoustics of sound propagation (Houtgast and Steeneken 1985, Rakerd and Hartmann 1986, Yost 2021), may limit the temporal precision with which auditory responses can be predicted. Given that input timing is often difficult to precisely predict and that hyperpolarization-activated conductances are common in neurons, cellular/synaptic mechanism of the type described here may be broadly relevant for predictive processing in the nervous system.

## Acknowledgments

This work was supported by a grant from the National Institutes of Health (NIDCD R01DC015449) to N.B.S.

## Author contributions

N.B.S., S.Z.M., Q.Z., and L.F.A. designed the research. Q.Z. collected the data. Q.Z., S.Z.M., and N.B.S. analyzed the data. S.Z.M and L.F.A. performed the computational modeling.

## Data and code availability

Data and code will be deposited on a public repository prior to publication.

## METHODS

### Animals

All experiments were conducted in accordance with National Institutes of Health (NIH) guidelines and with the approval of the Columbia University Institutional Animal Care and Use Committee. Adult male and female mice (C57BL/6J) aged 8-10 weeks were used for experiments. Mice were housed in an on-site animal facility on a 12 hr light-dark cycle. Experiments were performed during the light cycle. Calb1-Cre mice (Calb1-IRES2-Cre-D, stock# 028532) and floxed ChR2 mice (RCL-ChR2(H134R)/EYFP, Ai32, stock# 024109) were initially obtained from Jackson Labs.

### Surgery

Mice received subcutaneous injections of sustained release buprenorphine (0.75 mg/kg) before the surgery. Mice were subsequently anesthetized with isoflurane (1.5 ∼ 2%) and placed in a stereotaxic frame. The skull was exposed and a custom stainless steel headplate (5 x 25 x 1 mm) was attached to the skull with dental cement (C&B Meta-bond, Parkell, Edgewood, NY) such that the surface of the headplate was parallel with the horizontal plane connecting bregma and lambda. Mice were allowed to recover in their home cage for 3 days before the experiments. One day before the electrophysiology experiment, a 1.0 mm x 0.5 mm craniotomy was made above the DCN (around 6.0 mm posterior and 2.4 mm lateral to Bregma) on the right side and covered with silicone elastomer (Kwik-sil, World Precision Instruments, CA) until the time of the recording.

### Experimental apparatus and auditory stimulus presentation

All mouse behavior and neurophysiology experiments were performed in a double walled sound-attenuating chamber (Double Deluxe Model, Gretch-ken Industries). The ambient noise within the chamber was <30 dB SPL as measured by a sound pressure level meter (Bruel and Kjaer Type 2240). A customized headplate holder and mouse holding cup were positioned near the geometric center of the chamber. A food dispenser (ENV-203-20, Med Associates, VT) was placed 35 cm to the left of the mouse holding cup. The food pellets were delivered by the dispenser and transferred to the mouse by a plastic tubing (United States Plastic Corp, OH). A spoon-shape customized 3D-printed part was connected to the end of the plastic tubing to hold the food pellet.

Free field sound was presented via a Tucker-Davis Technologies (TDT, Alachua, FL) ES1 electrostatic speaker positioned ∼30 cm in front of the animal. The electrostatic speaker was covered with a grounded copper mesh to reduce electrical noise. Sound pressure levels of acoustic stimuli as measured in dB SPL were calibrated to the location of the animal’s right ear. The frequency response of the sound system was measured to be flat (+/−4dB) from 1 kHz to 50 kHz using a ¼’’ condenser microphone (377C01, PCP piezotronics), attached to a preamplifier (426B03, PCP piezotronics) positioned at the location of the mouse’s right ear. No attempt was made in this study to correct for the frequency response of the speaker output. Sounds caused by chewing were monitored by a small electret microphone (Knowles model 23329N). Microphone signals were sampled at 100 kHz and digitized using an analog to digital converter (Power 1401, Cambridge Electronic Design).

The waveforms of the acoustic stimuli were generated by customized MATLAB scripts with a 150 kHz sampling rate. The stimuli were 200 ms-long presentations of broadband noise band-pass filtered between 4 and 75 kHz or pure tones with 10 ms ramps at the beginning and end of the presentation. Rate level functions were determined using either broadband noise or characteristic frequency tones spanning from 5 to 75 dB SPL in 1 dB steps. The amplitudes of the sounds were randomly ordered during the presentation. Each stimulus was followed by a 400 ms-long interval. For characterizing the response map of the DCN units, pure tones ranging from 4 to 54 kHz were played at different amplitudes ranging from 5 dB SPL to 60 dB SPL. The frequencies were played in order from low to high.

### Behavioral training

Prior to recording, mice were food deprived (80-85% original weight) and habituated to eating food pellets (20 mg each, Bio-Serv, NJ) while head-restrained. Mice were considered habituated once they consumed more than 25 pellets within a 40-minute session for two consecutive days. Chewing behavior was filmed using a high-speed camera under IR illumination (CM3-U3-13Y3M-CS, Edmund Optics, NJ). Off-line video analysis used DeepLabCut to track the positions of the upper and lower jaw, and tongue during chewing behavior (Mathis, Mamidanna et al. 2018).

### Electrophysiology

Multi-site extracellular recordings were performed using 32-channel silicone probe electrodes (Cambridge Neurotech, UK). For optogenetic activation of CWCs, recordings were performed with an electrode fused to an optical fiber (H6b or H10b 32-channel probe + 100 µm core flat-end optical fiber 0.37NA, Cambridge NeuroTech, UK). Electrodes were connected to a headstage (Intan Technology) and data sampled at 30 kHz (Open Ephys, https://open-ephys.org/acq-board). DCN was targeted for recording using a vertical approach by slowly lowering the electrode vertically through the overlying cerebellum (5.85-6.25 mm posterior, 2.35-2.55mm lateral to Bregma) using a micromanipulator (IVM, Scientifica, UK). As the electrode was advanced through the cerebellum a series of 200 ms long search tones from 5 kHz to 50 kHz (in 5 kHz steps) or broadband noise were delivered. Entrance into DCN was marked by the sudden appearance of sound-evoked multi-unit activity which typically occurred ∼2500-3000 μm below the surface of the cerebellum. Food pellets were delivered to the mice during the recording session to characterize neural responses to self-generated sounds. Auditory stimuli were delivered in-between pellet consumption to characterize responses to external sounds and to aid in cell-type classification. Data collection was restricted to the DCN. Passage from DCN into VCN was determined by monitoring the tone frequency that most strongly drove multi-unit activity at sites located near the tip of the electrode. As the electrode advanced ventrally, the best frequency for driving multi-unit activity progressively decreased. A sudden increase in the best frequency (generally from ∼5 kHz to ∼20 kHz and usually occurring 600-700 μm below the surface of DCN) signified entrance into the VCN.

### Electrophysiological data analysis

Following the data acquisition, automated spike sorting was performed using Kilosort 4.0 followed by manual curation using Phy. Units accepted for subsequent analysis had high signal-to-noise ratios (mean 21.9 for FCs and 18.7 for CWCs), refractory period violation rate <0.5% (mean 0.13%), and a false negative rate <5% (mean 0.53% for FCs and 1.35% for CWCs). The latter was estimated by fitting each unit’s spike amplitude distribution with a Gaussian function and quantifying the fraction of spikes falling below the detection threshold (Hill, Mehta et al. 2011, Laboy-Juárez, Ahn et al. 2019, Fabre, Van Beest et al. 2023).

Units were identified as CWCs based on the presence of distinctive complex spike bursts. Complex spikes are stereotyped, high-frequency action potential bursts superimposed on a slower depolarization and are not observed in any DCN cell types except CWCs (Zhang and Oertel 1993, Manis, Spirou et al. 1994). Similar to previous *in vivo* extracellular recording studies of mouse (Portfors and Roberts 2007, Singla, Dempsey et al. 2017), we defined complex spikes as high-frequency bursts consisting of at least 3 spikes with interspike intervals between the first and second spike ≤ 4 ms and the interspike intervals between the second and third spike ≤ 3.5ms. Later spikes were also considered to be spikelets in a complex burst unless their interspike interval was ≥ 5 ms or their amplitude was larger than 80% of the first spike in the burst. Units were identified as FCs based on their responses to pure tones and noise and high spontaneous firing rate. Most putative FCs had type III or type IV response maps and spontaneous firing >20 Hz (Young and Brownell 1976, Hancock and Voigt 2002, Ma and Brenowitz 2012). Units with type II response maps and low or zero spontaneous firing were categorized as putative vertical cells and are not included in the present analysis (Rhode, Smith et al. 1983). Data from n = 46 mice were used for analysis of CWC and FC responses to auditory stimulation and chewing in **Figures 1-2** & **4**.

### Optogenetic experiments

CWC-specific ChR2 expression was achieved using the Calb1-Cre driver line (JAX #028532). To assess the specificity and coverage of Cre-mediated recombination within the DCN, Calb1-Cre mice were crossed with a fluorescent reporter line (Ai9; JAX #007909) and DCN sections were immunostained for PCP4, an established and selective marker for CWCs in the DCN (Wouterlood and Mugnaini 1984, Berrebi, Morgan et al. 1990). Of all PCP4-positive cells, 89% co-expressed the tdTomato reporter, and 92% of all reporter-positive cells within the DCN were PCP4-positive (Fig. 3A), indicating good coverage and specificity for CWCs. Although Calb1 is also expressed in Purkinje cells of the overlying cerebellum, expression within the DCN itself was largely restricted to CWCs. *In vivo* confirmation of opsin expression was obtained in each recording session by verifying that brief laser pulses reliably evoked complex spikes in a subset of recorded units, consistent with direct ChR2-mediated depolarization of CWCs. Optogenetic stimulation consisted of light pulses (10 ms duration, 1.0–2.0 mW at the fiber tip) delivered via a 470 nm laser (Coherent, OBIS) through an optical fiber integrated with the recording probe.

Chewing-optogenetic pairing. Jaw position was tracked in real time throughout the experiment using high-speed video (100 f.p.s.) and Bonsai software. The experiment consisted of three sequential phases. In the pre-pairing phase, mice consumed food pellets while baseline CWC and FC responses to chewing were recorded; no laser stimulation was delivered. In the pairing phase, mice consumed 20–25 pellets in sequence. The laser was triggered each time the lower jaw reached its lowest position (maximum opening), as detected online using Bonsai. Closed-loop triggering was active only during periods of regular rhythmic chewing and was suspended between chewing bouts to prevent spurious triggering by unrelated jaw movements. In the post-pairing phase, mice consumed pellets under the same conditions as the pre-pairing phase, again without laser stimulation. Experiments in which optogenetic stimulation was paired with BBN (**Fig. 3H-J**) consisted of alternating presentation of BBN alone (∼200 presentations) and BBN paired with optogenetic stimulation following a 30 ms delay (300-600 presentations). These two phases were alternated until approximately 2000 paired presentations were accumulated. Data from n = 8 ChR2 positive and n = 4 littermate controls are included in **Figure 3**.

### Immunohistochemistry and microscopy

Mice were anesthetized with an intraperitoneal injection of a ketamine (100 mg/kg) and xylazine (10 mg/kg) mixture and perfused with phosphate buffered saline (PBS, pH = 7.4, MRGF-6236, Growcells, CA), followed by a solution containing 4% paraformaldehyde in PBS. Brains were removed and postfixed in the same solution for 2 h before transferred to 30% sucrose (Sigma-Aldrich, MO) solution. Coronal sections of the DCN were cut at 50 microns on a cryostat. Slices were rinsed 3 times (10 min each) in PBS then incubated in a PBS solution containing triton-X (Sigma-Aldrich, MO), normal goat serum (0929391-CF, MP Biomedicals, CA) and primary antibodies (rabbit anti-PCP4, 1:500, Atlas Antibodies, HPA005792 ; guinea pig anti-tagRFP, 1:1000, CancerTools, 155266) overnight at 4 degrees. Slices were then rinsed 3 times for 10 minutes in PBS and incubated in a PBS solution containing Triton-X-100, normal goat serum and fluorescent secondary antibodies for 2 hours at room temperature, followed by 3 washes in PBS for 10 minutes. Slices were then mounted on slides (Superfrost Plus, Erie Scientific, 4951PLUS-001) and covered with the mounting medium (0100-01, SouthernBiotech) and a glass coverslip. Images were acquired by W1-Yokogawa inverted spinning disk confocal (Nikon, Japan). Images were processed using standard routines in Fiji (ImageJ).

### Modeling

The model consists of granule cell inputs and a CWC generating a simple spike rate and complex spikes. The total input to the CWC consists of granule cells firing in a delay line pattern and a strong auditory input from granule cells firing at sound onset. The granule cells (*g_i_*) generate EPSCs in the CWC through

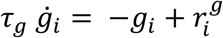

where *τ_g_* = 15 ms when we use the broader learning rule (**Figure 5**) and *τ_g_* = 5 ms when we use the narrower learning rule (**Figure S4**). The CWC simple spike rate is

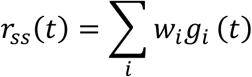

The simple spike rate is used as a stand-in for the CWC membrane potential to determine the T-type Ca^++^ current in the model, which is used to generate complex spikes. The dynamics of the T-type activation (*h*) is modeled as

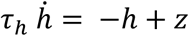

where z = 1 − 0.77*r_ss_*/*r_ss_^bl^* with *r_ss_^bl^* the baseline simple spike rate. This equation ensures that, at baseline, *h = 0.23*. We further bound *z* ∈ [−1,1] and ℎ ∈ [0,1]. *τ*_ℎ_ = 10 ms when we use a narrow learning rule. However, when using a broader learning rule the value of *τ*_ℎ_ depends on the value of *z*, whereby

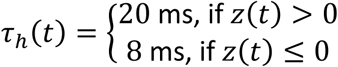

which means that *h* inactivation (when the cell is strongly depolarized and *z<0*) is faster than *h* activation (*z>0*) (Smith and Sherman 2002).

Complex spikes (CS) have a baseline firing rate of 1 Hz. In addition, the complex spikes rate is determined by the T-type Ca^++^ activation and the degree of the depolarization, as indicated by the simple spike rate (Smith and Sherman 2002),

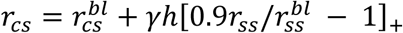

This implies that the depolarization threshold for evoking CSs is 1.1 times the baseline simple spike rate. []_+_indicates rectification so the complex spike rate does not go below 0. *γ* = 75 Hz when we use the narrow learning rule and *γ* = 200 Hz when we use the broader learning rule.

Plasticity at g-cell-CWC synapses is driven by the following rule:

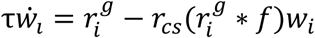

where τ is the learning rate and *f* is the kernel that defines the eligibility trace (i.e. *f* is the learning rule window). We made the second term here multiplicative to ensure stability and initialize all *w* equal to 1. For the narrow learning rule window, we model *f* as a difference of exponentials,

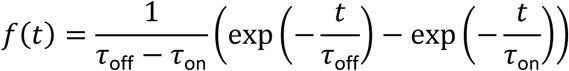

where *τ*_off_ = 5 ms and *τ*_on_ = 3 ms. For the broader learning rule window, we model *f* as,

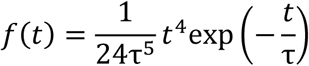

where τ = 12 ms

To prevent learning from depressing the g-cell input that drives the CS when we use the narrow learning rule window, we set the inputs driving auditory response to be non-plastic (as described in the text).

## SUPPLEMENTARY FIGURES

**Figure S1.**
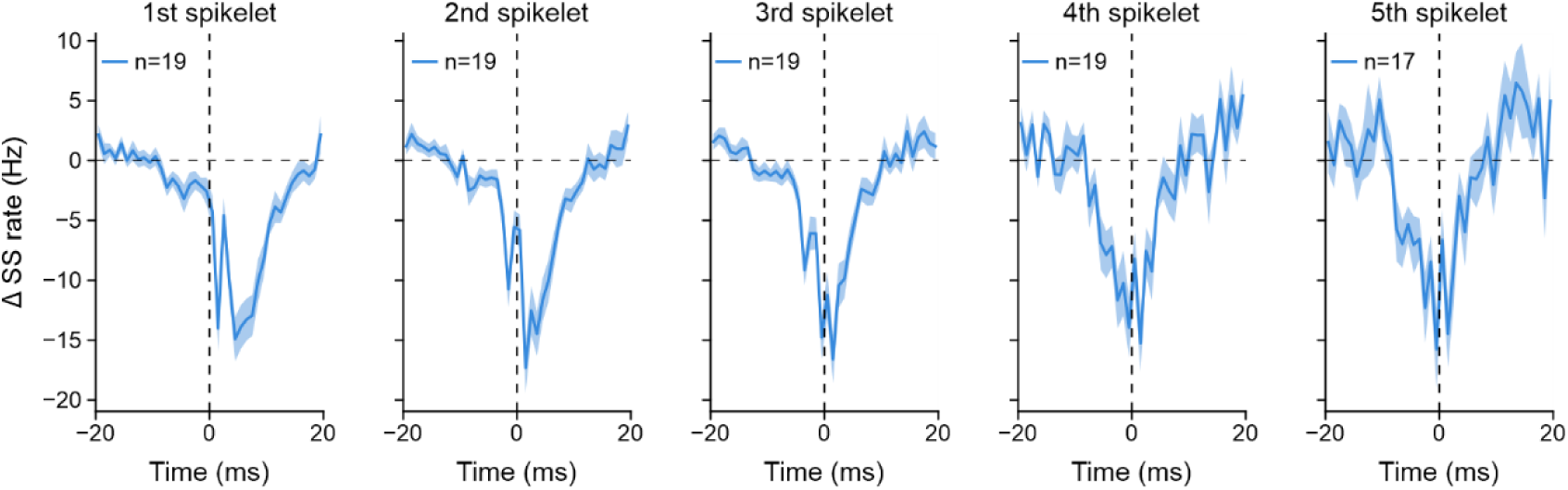
Each spikelet of the complex spike contributes to CWC-mediated inhibition of FCs. Average FC spike rate aligned to the time of each successive spikelet within complex spike bursts in simultaneously recorded, putatively monosynaptically connected CWC-FC pairs (n = 19 pairs). Each spikelet is associated with a transient reduction in FC firing rate of similar magnitude, consistent with faithful axonal propagation of high-frequency spike bursts in CWCs and reliable transmission at CWC-FC inhibitory synapses *in vivo*. Related to Figure 1.

**Figure S2.**
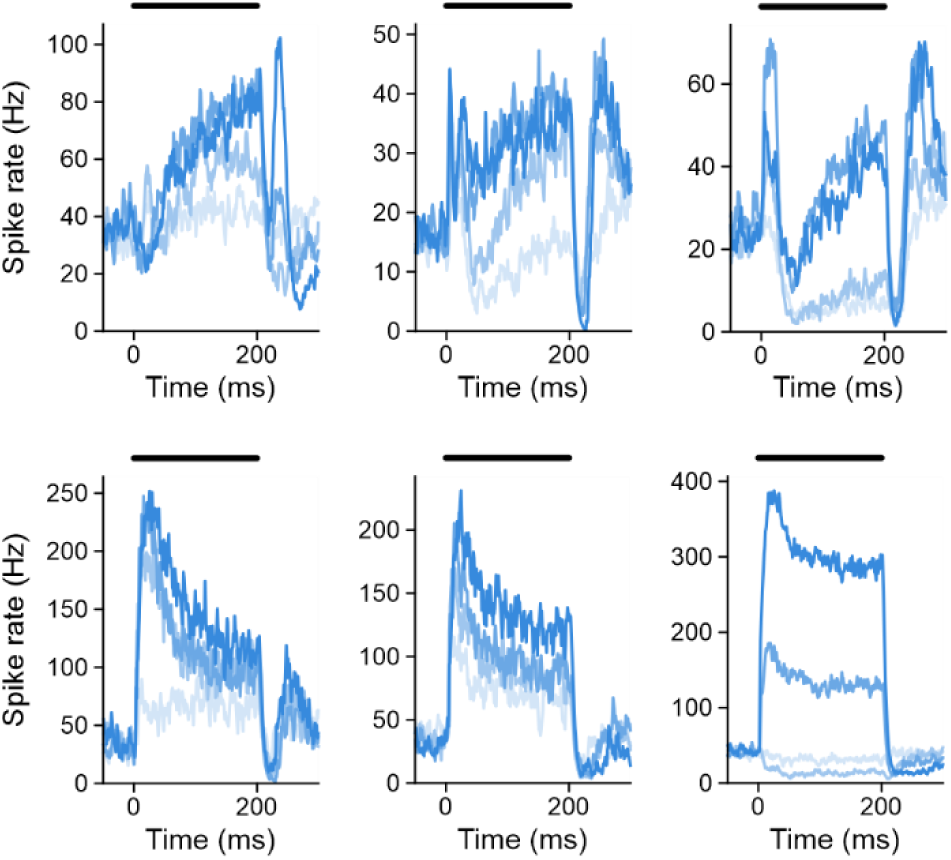
Heterogeneous temporal response profiles in FCs. Responses to BBN across a range of sound levels for 6 example FCs. Temporal response profiles varied across units and as a function of sound level. Sound levels were 5-25, 25-40, 40-60, and 60-75 dB SPL from lightest to darkest. Related to Figure 1.

**Figure S3.**
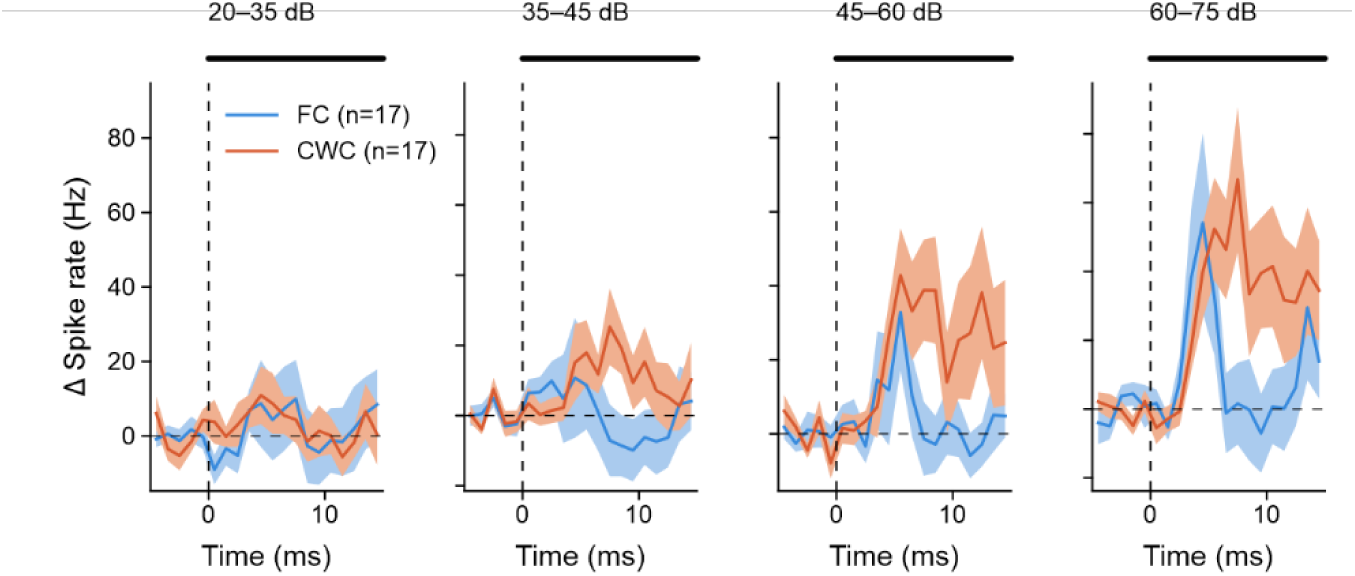
Matching temporal profiles of auditory response in synaptically-coupled CWCs and FCs. Population average responses of putative synaptically-coupled FCs (n = 17) and CWCs (n =17) to the onset of BBN across sound levels. CWC spike rate includes simple spikes, complex spikes, and spikelets. Related to Figure 1.

**Figure S4.**
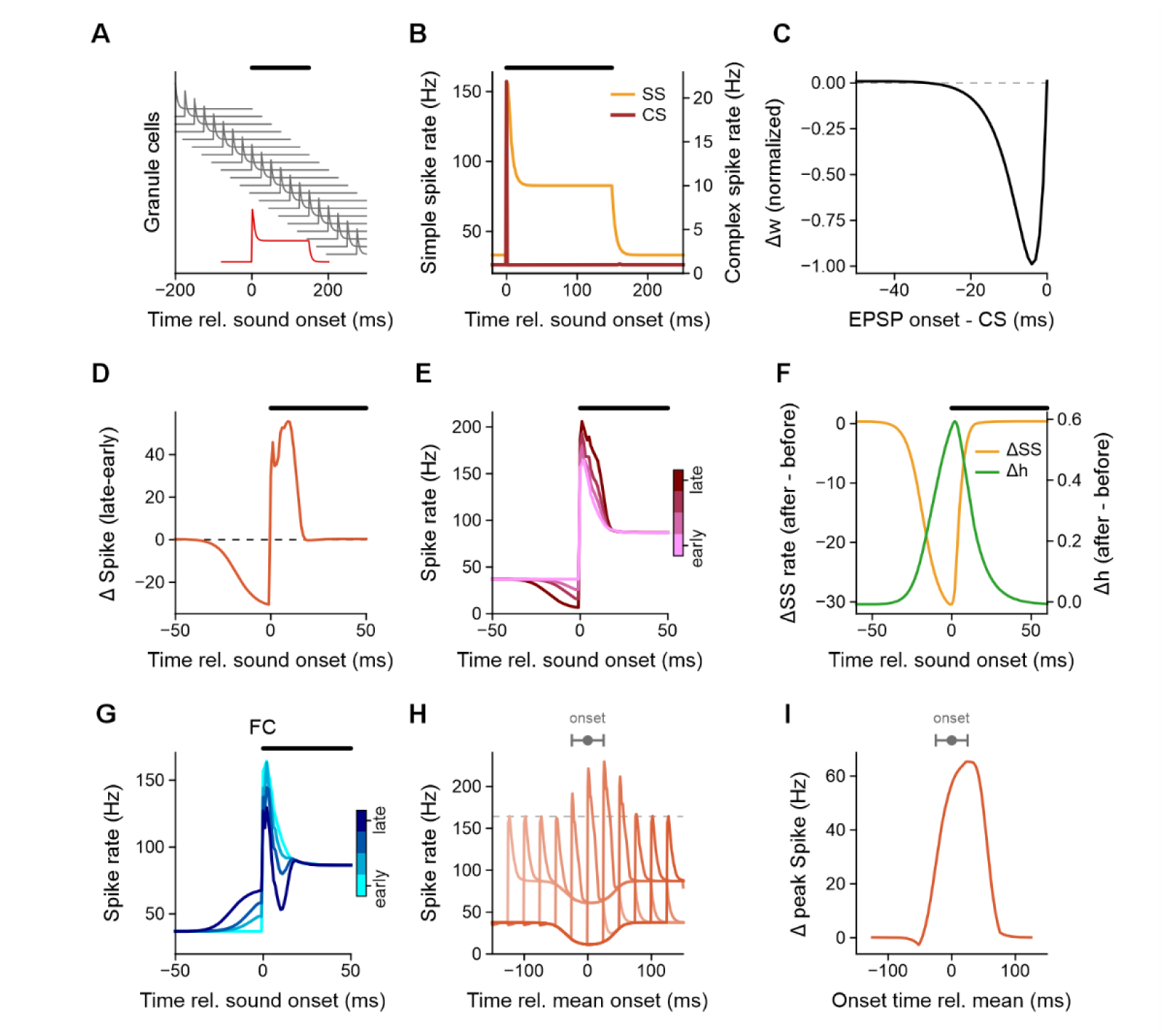
Simulations with a narrower anti-Hebbian learning rule window and non-plastic auditory granule cell input also reproduces the positive-feedback mechanism that enhances CWC responses to predicted sound onset. **(A)** Schematic of model inputs to a CWC. Predictive granule cell inputs (grey) tile the time surrounding the sound, whereas auditory inputs (red) provide excitation beginning at sound onset. Auditory granule cell inputs are assumed to be non-plastic in this version of the model (see **Discussion**). The model also includes inhibition simulating superficial stellate cells (not shown). **(B)** Model CWC response before learning. Auditory input drives a transient increase in simple spike (SS) and complex spike (CS) rates at sound onset, followed by a sustained increase in simple spike rate. **(C)** Anti-Hebbian plasticity rule used in this version of the model. Associative depression is restricted to a narrow temporal window, matching *in vitro* data. (**D-I**) These plots replicate the results shown in Figure 5 but for this version of the model using a narrower learning rule window and non-plastic auditory granule cell input. Related to Figure 5.

